# Predictive Coding of Novel versus Familiar Stimuli in the Primary Visual Cortex

**DOI:** 10.1101/197608

**Authors:** Jan Homann, Sue Ann Koay, Alistair M. Glidden, David W. Tank, Michael J. Berry

## Abstract

To explore theories of predictive coding, we presented mice with repeated sequences of images with novel images sparsely substituted. Under these conditions, mice could be rapidly trained to lick in response to a novel image, demonstrating a high level of performance on the first day of testing. Using 2-photon calcium imaging to record from layer 2/3 neurons in the primary visual cortex, we found that novel images evoked excess activity in the majority of neurons. When a new stimulus sequence was repeatedly presented, a majority of neurons had similarly elevated activity for the first few presentations, which then decayed to almost zero activity. The decay time of these transient responses was not fixed, but instead scaled with the length of the stimulus sequence. However, at the same time, we also found a small fraction of the neurons within the population (∼2%) that continued to respond strongly and periodically to the repeated stimulus. Decoding analysis demonstrated that both the transient and sustained responses encoded information about stimulus identity. We conclude that the layer 2/3 population uses a two-channel predictive code: a dense transient code for novel stimuli and a sparse sustained code for familiar stimuli. These results extend and unify existing theories about the nature of predictive neural codes.

## Introduction

Adaptive and coordinated behavior requires that an animal be able to make predictions about the near and far future. This intuition that some neural computations should be ‘predictive’ has a long history, starting with ideas about how the receptive field structure of retinal ganglion cells relates to the statistics of natural visual scenes (Atick and Redlich, 1990, 1992; Attneave, 1954; Srinivasan et al., 1982). More recently, it has been discovered that other retinal circuits carry out more active forms of prediction about the visual world, including predictions about the future location of objects moving at constant velocity (Berry et al., 1999; Bullock et al., 1990; Johnston and Lagnado, 2015; Schwartz et al., 2007b; Trenholm et al., 2013) and about temporally periodic sequences of light intensity (Schwartz and Berry, 2008; Schwartz et al., 2007a). Through these computations, the retina composes a population neural code that represents visual information that was *predictable* to the retina separately from information that was *surprising* (Berry and Schwartz, 2011) and can efficiently capture the predictive components of the overall sensory information (Palmer et al., 2015). Thus, some of the diversity of retinal circuitry is used to make predictions about the upcoming visual stimulus, and some of the redundancy of the population code is used to represent visual stimuli that are predictable as well as those that are surprising.

Given the importance of predictive computations for an animal’s behavior, one might expect that other circuits in the brain perform additional predictive computations, not performed by the retina. In particular, related ideas about predictive computation have also been influential in theories about the function of the neocortex (Barlow, 1994; Bastos et al., 2012; Heeger, 2017; Mumford, 1992; Rao and Ballard, 1999). Here, the relatively stereotyped local circuitry of the neocortex has long led to speculation that each local circuit might be carrying out a somewhat similar, fundamental computation on its specific inputs. In addition, the organization of sensory-motor pathways into hierarchies (e.g., V1 → V2 → V4 → IT in the ventral visual stream) with stereotyped feedforward and feedback connections (Felleman and Van Essen, 1991; Markov et al., 2014; Rockland and Pandya, 1979) has motivated ideas about hierarchical predictive codes, where higher levels of the hierarchy send predictions down to lower levels of the hierarchy that then compare their inputs against those predictions and only send the surprises up the hierarchy (Bastos et al., 2012; Mumford, 1992; Rao and Ballard, 1999). On the other hand, this same anatomy has motivated another theory in which predictions are fed forward in the hierarchy and errors are fed back (Heeger, 2017). The experiments reported here were motivated by the broad hypothesis, based on this prior work, that predictive computations might be distributed throughout the sensory hierarchy, and that they might constitute an important piece of the core set of computations carried out in each local neocortical circuit.

Among the simplest kinds of predictable sensory patterns are periodic stimuli. When presented with a violation of an ongoing periodic stimulus, many areas of the cortex generate a specific response to that violation. Examples include the mismatch negativity (MMN) (Naatanen et al., 2007; Naatanen et al., 1982) as well as analogs in other sensory cortices (Bullock et al., 1994; Klinke et al., 1968; Sutton et al., 1967) observed in human EEG and in Ca^++^ imaging data (Hamm and Yuste, 2016). On a cellular level, stimulus-specific adaptation (SSA) enhances the response of neuron in the primary auditory cortex to a rare stimulus (Ulanovsky et al., 2003), an effect that is further enhanced when the rare stimulus violates a periodic pattern (Yaron et al., 2012). Similarly, neuronsin the primary visual cortex exhibit suppression to common stimuli and enhancement to rare stimuli (Hamm and Yuste, 2016; Vinken et al., 2017). When a natural movie clip is repeated many times, responses become more reliable and sequences of activity subtly reverberate when the stimulus ends (Yao et al., 2007). Similarly, when the cortex is trained with a spatial sequence of activation, stimulation at the initial spatial location triggers a subtle reactivation of the whole sequence (Xu et al., 2012). On a longer time scale, repetition of the same stimulus sequence over multiple days enhances the neural response (Frenkel et al., 2006) and leads to pattern completion of a missing image (Gavornik and Bear, 2014). Finally, a mismatch between expected versus actual visual stimuli during closed-loop running can generate a strong response in V1 (Keller et al., 2012; Zmarz and Keller, 2016).

Here, we investigate predictive processing of repeated stimuli in the primary visual cortex. This cortical area is advantageous because it is the first stage in the cortical visual hierarchy, so its inputs embody relatively few prior computations. We formed temporal sequences from images having a random assortment of line segments, and presented the same temporal sequence repeatedly to an awake, head fixed mouse. While this repeated temporal sequence was ongoing, we occasionally substituted images with novel images drawn from the same distribution. We first trained mice to lick in response to the presentation of a novel image. Animals readily learned this task, achieving significant performance on the first day of testing and generalizing to sequences of up to 7 images. Then, in a seperate set of experiments, we used two-photon Ca^++\^ fluorescence imaging to measure neural activity in layer 2/3 (Dombeck et al., 2010). We found that the majority of neurons exhibited excess activity in response to a novel image. Similarly, when we began presenting a new temporal sequence, a majority of the neurons exhibited elevated firing that relaxed to a steady-state after ∼2 presentations of the same temporal sequence. Interestingly, the dynamics of this adaptation process were not constant in time, but instead scaled with the duration of the temporal sequence. In addition, a small fraction of cells (∼2%) exhibited a strong periodic response to a given temporal sequence. When we changed the temporal sequence, a different sparse subset of neurons exhibited periodic responses. Finally, decoding analysis showed that both the transient and periodic responses conveyed information about the identity of the temporal sequence. Thus, the population code in layer 2/3 of primary visual cortex exhibits two channels of information: a sparse channel of highly active cells representing familiar or predictable stimuli, and a dense channel of weakly active cells for novel or surprising stimuli.

## Results

In order to explore how the visual system processes novel versus familiar images, we first tested whether mice could discriminate a novel image from otherwise similar repeated images. Because we wanted to focus on the role of early visual processing, we designed our stimuli to drive neurons in the primary visual cortex. We chose images that consisted of a superposition of Gabor patches (see Methods), which resemble a random assortment of line segments (Fig. 1A). We randomly generated a set of images from this ensemble and composed them into a temporal sequence, which was repeated many times (Fig. 1B; blue). In addition, we replaced a small fraction of these images with a novel image (Fig. 1B; orange). Because all images were drawn from the same ensemble, they were closely matched for low-level image statistics, like overall light level, contrast, and spatial scale (see Methods). Each image was displayed for 200 – 300 ms, a time long enough that we can rule out a retinal origin for any predictive processing (Schwartz et al., 2007a). In order to vary the level of difficulty of the task, we formed sequences containing from 2 up to 7 images.

**Figure 1.**
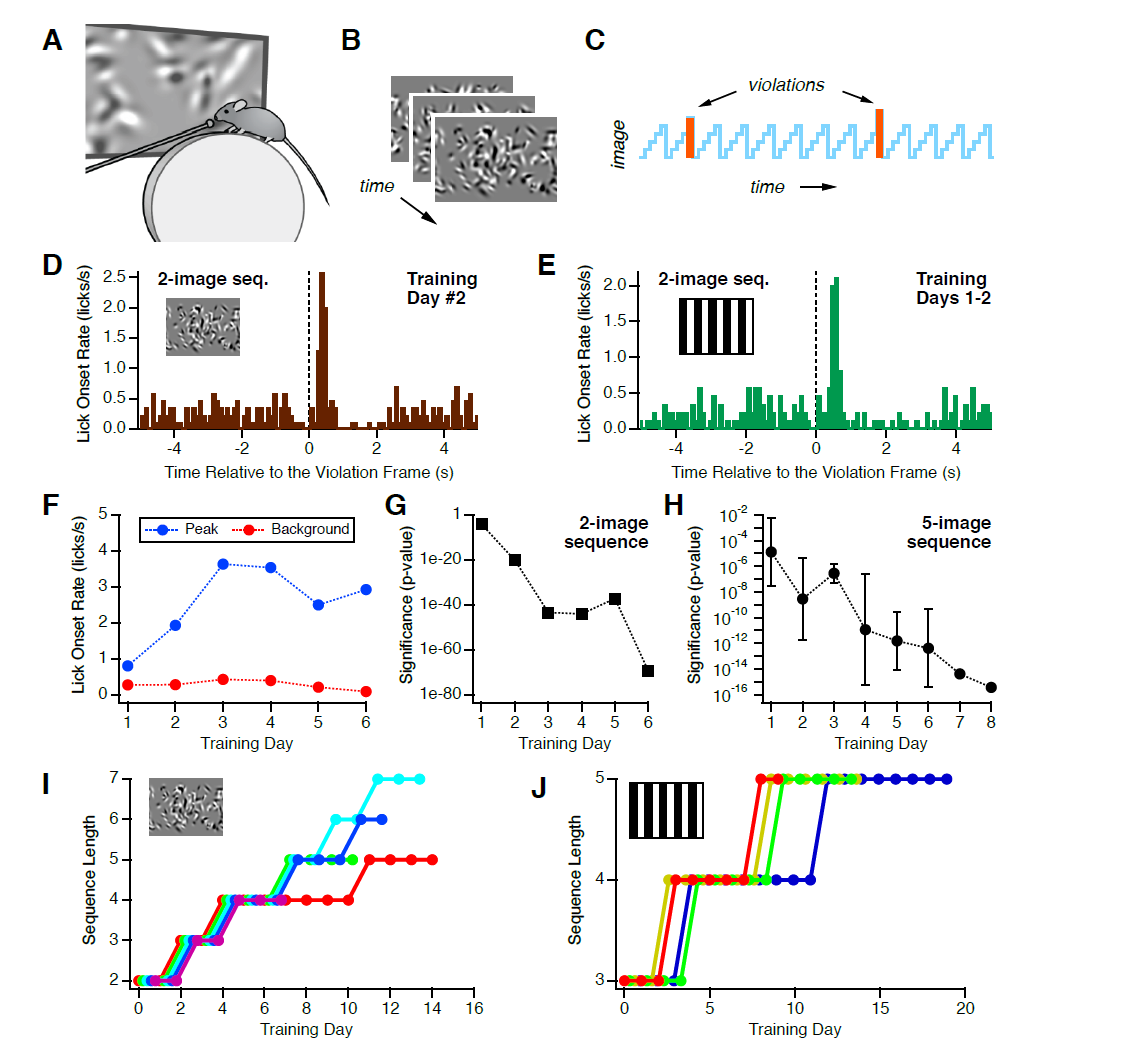
Mice can learn to lick in response to a novel image. **A.** Water restricted, head-fixed mice received water through a lickport while viewing stimuli. **B.** Temporal sequences were formed from several different images, each consisting of 100 randomly chosen, superimposed Gabor functions. **C.** Such a temporal sequence was repeated with novel images randomly inserted (orange). **D, E.** The rate of lick onsets as a function of time relative to the occurrence of a novel image for sequences formed from (Gabor, grating) images. **F.** Background (red) and peak (blue) lick onset rate as a function of number of training days with a 2-image Gabor sequence. **G.** Significance of the peak in the lick onset rate as a function of number of training days (1 mouse; 2-image Gabor sequence). **H.** Significance of the peak in the lick onset rate as a function of number of days of training with a 5-image Gabor sequence; averaged over 5 mice. **I, J.** Sequence length achieved at criterion performance (see Methods) versus number of training days for (Gabor, grating) sequences; colors are for different mice.

### Animal Behavior

Water restricted mice were head-fixed but free to run on a stationary, vertical Styrofoam wheel while they viewed the above described visual stimuli (Fig. 1A). A lick port made water available to animals when a novel image was presented. Under these conditions, animals engaged in a strategy of exploratory licking. After a 3-day shaping period, the rate of lick onsets (which is the first lick within a burst of licks) showed a sharp increase following the onset of a novel image (Fig. 1D). This elevated lick onset rate following a novel image is evidence that the animal learned to recognize the novel image and used this recognition to drive its licking behavior. Over six days of this training regimen, the peak lick onset rate rose and the background lick onset rate declined (Fig. 1F), resulting in a marked increase in statistical significance of the peak (Fig. 1G). Notice that the licking behavior was significant on the *first day* of training (probability of that the observed increase in lick onset rate at the time of a novel image was due to chance was p = 8.1e-5). This rapid acquisition of the task suggests that recognition of novelty, even in the case of abstract images, may be a fairly natural task for the animal.

In order to explore the generality of this behavior as well as to make stronger contact to the V1 literature, we repeated our paradigm using temporal sequences composed of full field static gratings having different orientations (see Methods). Again, we found that animals began to lick with a short latency after the onset of a novel image (Fig. 1E) and acquired this ability on the first day of training.

We next trained our mice to perform this task for longer temporal sequences. Our method was to add one new image to the previously repeated temporal sequence and train on the increased sequence length until the animal achieved sufficiently high performance. As for the 2-image sequence, the significance of the peak in lick onset rate at the time of the novel image steadily increased as a function of days of training (Fig. 1H). By this method, we were able to train animals to achieve highly significant behavioral performance for sequences of 5–7 Gabor images (Fig. 1I) and 3–5 grating images (Fig. 1J). As we only trained for ∼1 week in a given condition, it is possible that mice could recognize novel images in even longer temporal sequences with more training.

### Neurophysiology

We next explored the responses of neurons in the primary visual cortex of naïve mice under the same visual conditions. Our approach has been to use two-photon Ca^++^ fluorescence imaging in mice that were awake and head-fixed but free to move their legs on a styrofoam ball placed below them (Fig. 2A) (Dombeck et al., 2007; Harvey et al., 2012). Animals were from a *Thy1* line (GP5.3, Janelia) that expressed the protein GCaMP6f (Chen et al., 2013) in excitatory neurons. In order to identify the primary visual cortex, we first carried out large-scale brain mapping using drifting bars (Fig. 2B; see Methods). Then, we selected a field of view in V1 and imaged at cellular resolution in layer 2/3. We used custom software to identify regions-of-interest corresponding to cell bodies having a “halo” pattern of fluorescence indicating expression mostly in the cytoplasm (Fig. 2C; see Methods). The time course of the fractional change in fluorescence in a single ROI (ΔF/F) exhibited sparse events on a background (Fig. 2D). Because we were interested in the capacity of the visual cortex to carry out unsupervised learning of temporal sequences, we did not present water rewards during our neurophysiology experiments.

**Figure 2.**
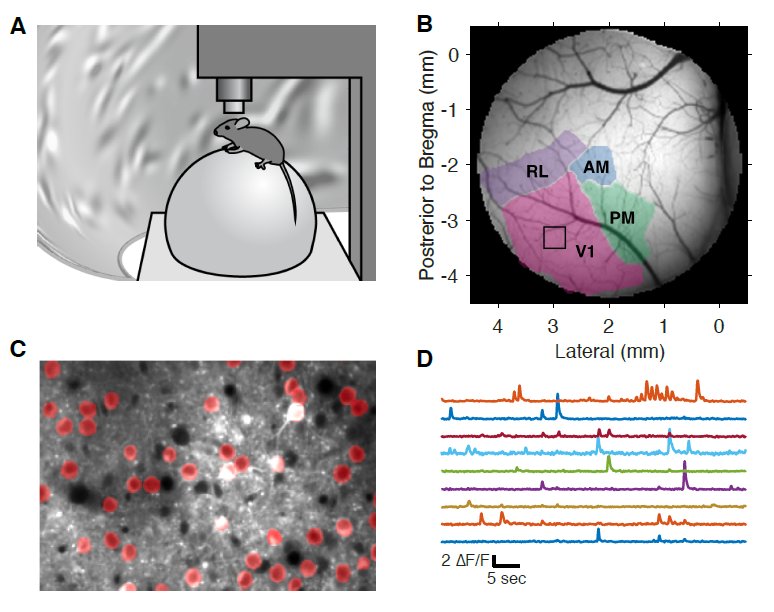
Two-photon calcium imaging. **A.** Head-fixed mice were awake and free to run on a Styrofoam ball, while visual stimuli were presented on a toroidal screen and neural activity was imaged through a two-photon microscope. Animals viewed the visual stimuli passively and performed no behavioral tasks. **B.** Results from brain area mapping, demarcating area V1 and showing the field-of-view for cellular resolution imaging (box). **C.** Field-of-view with identified regions of interest (ROIs; red). **D.** Example trace of neural activity in one ROI.

In the *Novelty Experiment*, we repeated a 4-image sequence with images of 250 ms duration. A single sequence was repeated for ∼10 min while novel images were randomly substituted every ∼5 sec (see Methods). Most neurons had a weakly modulated response to the repeated temporal sequence (Fig. 3A blue). But when a novel image was substituted, there was a large, transient response (Fig. 3A red). To quantify this effect, we subtracted the activity triggered on the repeated temporal sequence from that triggered on the presentation of a novel image to get the excess activity (Fig. 3A black). Within a population of over 1100 neural responses, the clear majority exhibited excess activity in response to novel images (Fig 3B; p<0.05 for 878/1134 = 77%, see Methods).

**Figure 3.**
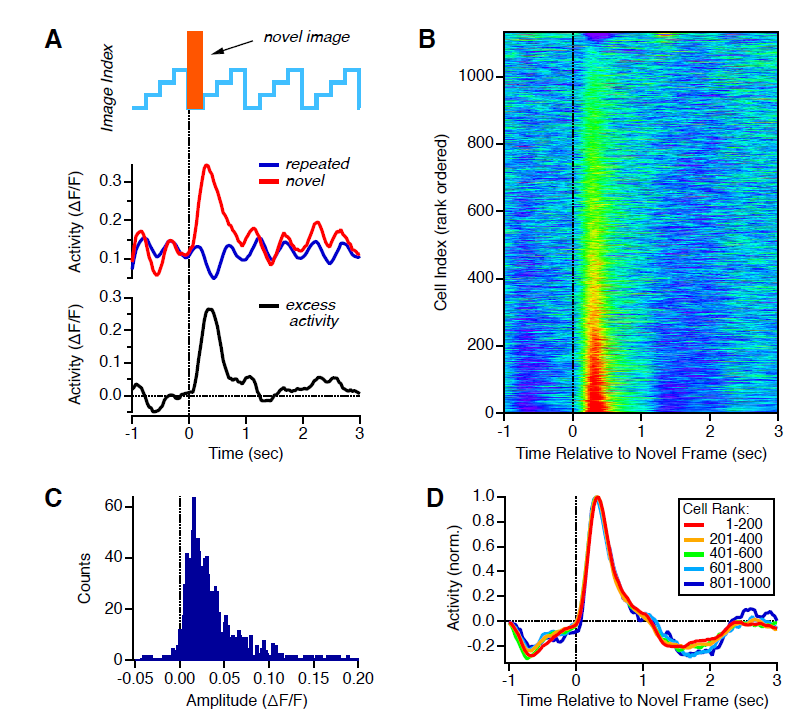
The response to a novel image. **A.** *Top*: Repeated temporal sequence containing a novel image. *Middle*: Activity of one example neuron with and without a novel image (red vs. blue). *Bottom*: Excess activity due to the occurrence of a novel image (black). **B.** Excess activity for 1134 neural responses (rows) plotted versus time relative to the occurrence of a novel image (color scale = z-score). **C.** Histogram of amplitudes of the excess activity. **D.** Activity versus time for different, rank-ordered groups of neurons (colors).

The amplitude of this novelty response varied in the population (Fig. 3C), ranging up to ΔF/F ∼ 0.2. When we converted this amplitude into a rough estimate of the number of spikes, we found that the number of excess spikes was ∼0.5 spikes per neuron per novel image (Supplemental Fig. S1). While this response may seem small, it is important to note that this analysis averages over many different examples of novel images. In other words, the number of excess spikes elicited by a single novel image within layer 2/3 of the primary visual cortex is ∼150,000 (see Methods) – a signal that could very well be salient to downstream visual areas (see Methods). We found that the latency and temporal dynamics of the novelty response were independent of the amplitude (Fig. 3D). These results were substantially unaffected by a maximal choice of neuropil subtraction (Supplemental Fig. S2). At the same time, the neuropil itself showed clear excess activity after the novel image (Supplemental Fig. S3), which is consistent with the fact that a majority of measured neurons also showed a novelty response.

It is known that Ca^++^ signals in V1 can be modulated by the animal’s locomotion (Keller et al., 2012; Niell and Stryker, 2010; Saleem et al., 2013). Therefore, we averaged the Ca^++^ response to a novel image over periods when the animal was either running or still. We did not find significant differences (Supplemental Fig. S4B,D). In addition, sudden unexpected visual stimuli, like dark looming objects from above (Yilmaz and Meister, 2013) or the onset of a bright light (Godsil and Fanselow, 2004) can trigger a defensive response, causing either flight or freezing. We therefore monitored running speed during our experiments and correlated speed with the onset of a novelty response. We did not observe a change in running speed at the onset of a novel image (Supplemental Fig. S4C,E). From these control analyses, we conclude that the novelty response is not the result of the animal's locomotion or of a defensive response to the visual stimulus.

Exposure to stressful stimuli leads to an activation of the sympathetic nervous system, which has been shown to dilate the pupil through the neural pathway of amygdala → locus coeruleus → Edinger-Westphal nucleus → pupil (see (Samuels and Szabadi, 2008) for a comprehensive review). Perhaps a novel image in our experiment evokes this form of stress, analogous to a ‘startle response’ evoked by a loud auditory stimulus? To address this possibility, we tested whether the presentation of a novel image caused a change in the pupil diameter. It did not (Supplemental Fig. S5C). Similar to running, pupil dilation increased the background neural activity but did not qualitatively change the novelty response (Supplemental Fig. S5B,D,E). In fact, pupil diameter correlated with running (Supplemental Fig. S5F) (McGinley et al., 2015). We also tested whether the novel image triggered saccadic eye movements. Again, it did not (Supplemental Figs. S6, S7), nor did periods with many eye movements qualitatively change the novelty response. Together, these analyses strongly suggest that the novelty response should not be interpreted as a form of generalized startle response. While this result might seem counter-intuitive, it is consistent with the fact that the presentation of a novel image under these conditions has very low salience for human observers. Readers are encouraged to view video clips of the visual stimulus (Supplemental Videos).

Conceptually, a neuron cannot exhibit a novelty response until the underlying temporal sequence is repeated at least once. Thus, we wanted to characterize the dynamics over which this response emerges. To this end, we designed a *Variable Repetition Experiment* in which we displayed a set of 3-image sequences of 300 ms per image, denoted by *(ABC)*_*i*_, where *i* is the sequence index (Fig. 4A; see Methods). Each sequence repeated for *L*_*i*_ times before a novel image was displayed (and which was followed by one more repeat to help us distinguish responses to the novel image from responses to the next sequence), together forming an 'adaptation block'. Different values of *L*_*i*_ taken from the set [2, 4, 9, 19, 39] were randomly interleaved, allowing us to vary the number of repeats of the same temporal sequence (which we call 'cycles') before the presentation of a novel image. Each sequence was unique.

**Figure 4.**
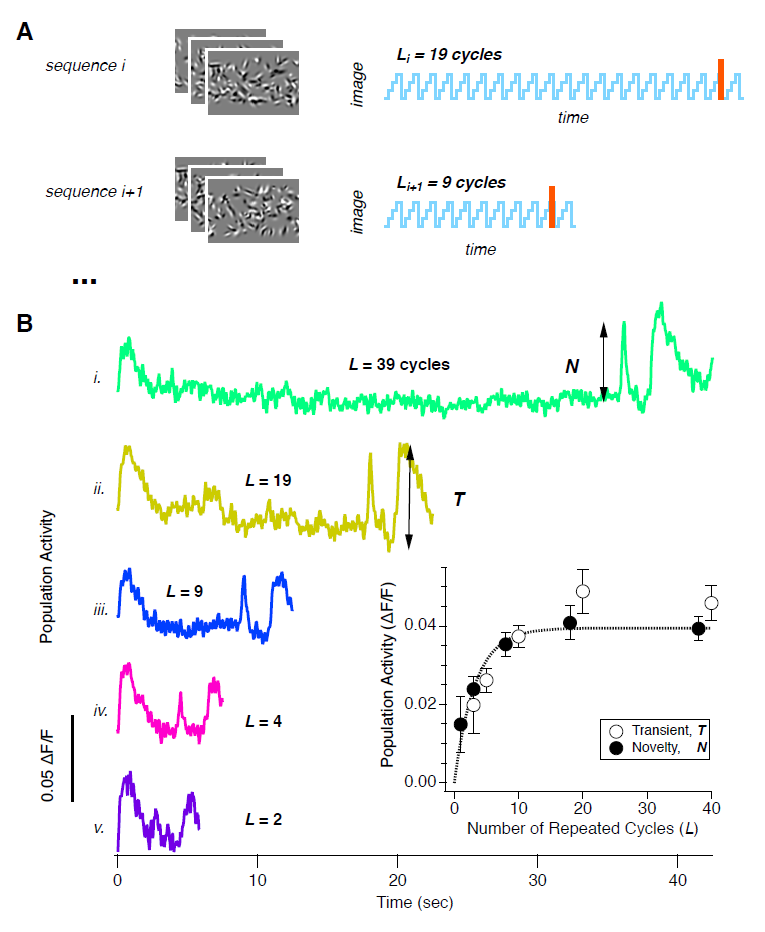
Dynamics for the Emergence of the Novelty Response. **A.** In the *Variable Repetition Experiment*, each adaptation block *i* had a new temporal sequence along with a random choice of the number of cycles of that sequence before a novel image, *L*_*i*_. Each block then had one more cycle of the temporal sequence. Then a new block began without blanks or interruptions. **B.** Neural activity averaged over the entire population for different choices of *L* (curves offset for clarity). *Inset*: Average activity for the novelty response (filled circles) and the transient response (open circles) plotted as a function the number of repeated cycles, *L*, of the temporal sequence along with an exponential curve fit constrained to run through the origin (dashed line); error bars are standard error over n=5 mice.

We found that novelty responses emerged rapidly. Significant excess activity was found in the population after as few as 1 repeated cycle (Fig. 4B). The effect increased with more cycles and saturated after ∼20 cycles (Fig. 4B inset). Fitting an exponential curve to the effect amplitude versus number of cycles revealed a time constant of 3.2 ± 0.7 cycles, or alternatively 2.9 ± 0.7 sec, for the emergence of the novelty response.

### The Transient Response

We also noticed that neurons exhibited elevated activity when we began presenting a new temporal sequence. In this experiment, each adaptation block was always preceded by a block that used a different sequence. Therefore, this initial elevated activity is conceptually very similar to the novelty response. We quantified the emergence of this transient response by measuring its amplitude as a function of the number of cycles, Li-1, in the previous adaptation block (Fig. 4B). This amplitude agreed closely with the amplitude of the novelty response and exhibited the same time scale of emergence (Fig. 4B inset). This result is expected from causality, because both a novel image and the first image of a new temporal sequence violate the periodicity of an ongoing temporal sequence. Of course, a new temporal sequence presents several novel images in a row, and as a result, the response is more extended in time than is the response to a single novel image.

In addition, the transient response adapted strongly as a given temporal sequence was repeated, quickly reaching a steady-state (Fig. 4B, elevated activity near t=0). This adaptation process had a time constant of 1.4 ± 0.4 sec in the *Variable Repetition Experiment*. This time scale was significantly shorter than that for the emergence of the transient response (p < 3•10^-7^), suggesting that different circuit mechanisms may be involved.

We next wondered whether the time scale of adaptation was fixed, as one might expect from a single biophysical mechanism, or variable, depending on the properties of the temporal sequence. To this end, we designed a *Variable Sequence Length Experiment*, in which we varied the sequence length. Similar to the *Variable Repetition Experiment*, we formed adaptation blocks. In each block *i*, we randomly chose the number of images in the sequence, *S*_*i*_, from the set [3, 6, 9, 12] and then randomly generated a new temporal sequence of this length (Fig. 5A). All images had the same duration, 300 ms, so that the total sequence duration ranged from 0.9 sec up to 3.6 sec. In all adaptation blocks, the given sequence was repeated for 20 cycles.

**Figure 5.**
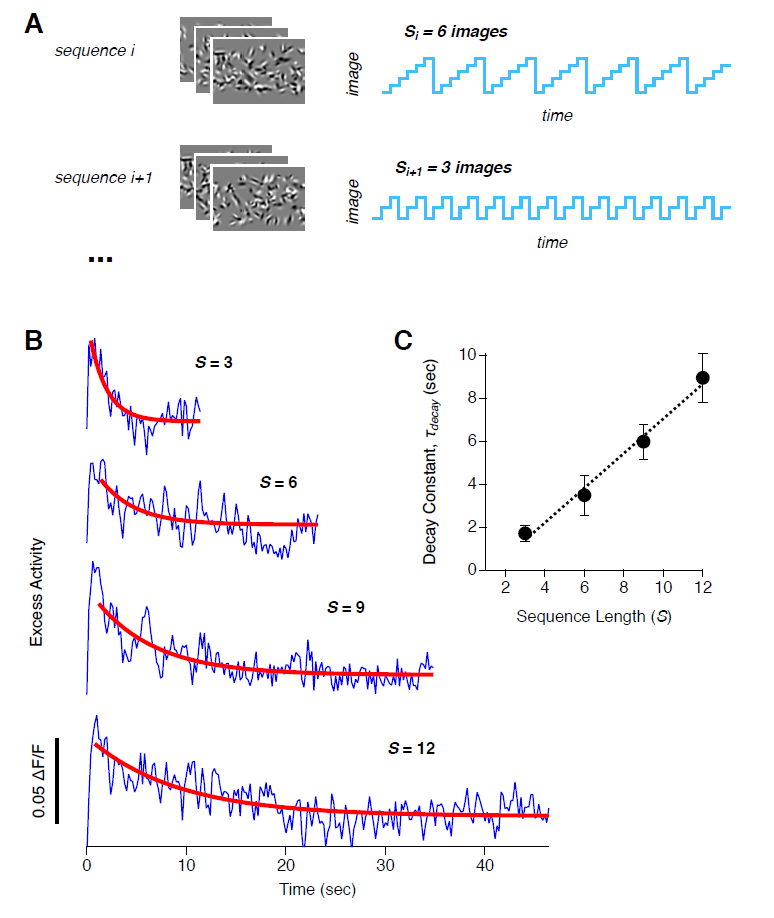
Flexible Dynamics of Adaptation to a Repeated Temporal Sequence. **A.** In the *Variable Sequence Length Experiment*, each adaptation block *i* had a new temporal sequence along with a random choice of the sequence length, Si. Each sequence was repeated for 20 cycles, and then a new block began without blanks or interruptions. **B.** Population average activity plotted versus time for sequences of different duration (blue), along with exponential curve fits (red); curves offset for clarity. **C.** Time constant of adaptation, decay, plotted against the sequence length, S.

We observed that the transient response decayed with slower dynamics when we repeated longer temporal sequences (Fig. 5B). Fitting an exponential curve to these dynamics, we found that the time constant of adaptation, *τ*_*decay*_, scaled linearly with the total temporal duration of the sequence (Fig. 5C). In other words, the dynamics of adaptation was constant in units of the number of sequence repeats, *τ*_*decay*_ = 2.1 ± 0.3 cycles.

### The Sustained Response

In order to gain more insight into these phenomena, we next examined the response across the entire population of neurons for a single choice of temporal sequence. In order to better average over neural noise, we designed a *Repeated Sequence Experiment*. This experiment also had adaptation blocks, but the difference was that we chose 10 different temporal sequences and repeated each of those blocks a total of 18 times (blocks for each sequence were randomly interleaved; see Methods). As before, we observed adaptation to repeated temporal sequences and transient responses to novel images. But in addition, a small fraction (roughly 2% under our visual conditions) showed a qualitatively different response type: a sustained response to the repeated stimulus (Fig. 6A,B, see the pink stars). This response was already present for the first presentation of the temporal sequence, consistent with feedforward models when part of the image matches a cell's receptive field. Importantly, when we displayed different temporal sequences, *different* neurons showed a sustained response (Fig. 6A vs 6B). This observation implies that sustained responses do not constitute a separate cell class, but instead are response types that are present in the same pool of neurons that can also show transient activity, depending on the identity of the temporal sequence.

**Figure 6.**
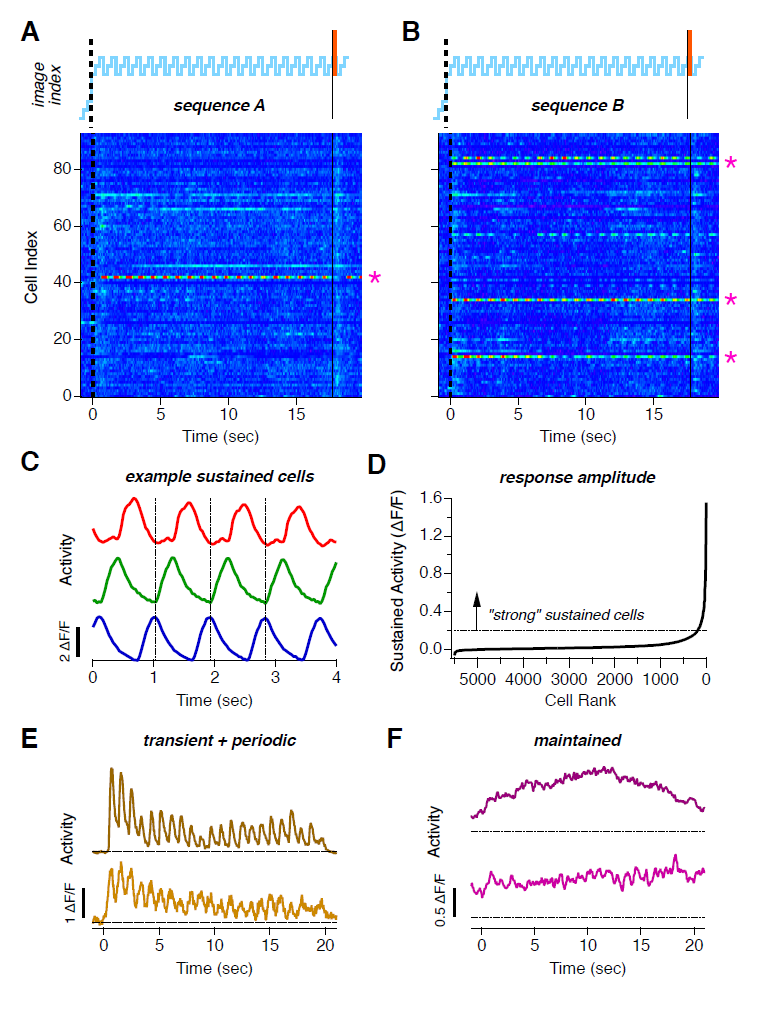
Sustained Responses. **A,B.** Matrix of activity of a population of the same neurons (n=93) responding to sequence (A,B); a given sequence starts at t=0 sec, novel image occurs at t=17.7 sec. Activity is averaged over 18 blocks with the same temporal sequence. Pink stars indicate neurons with periodic sustained responses. **C.** Activity versus time for 3 example neurons with sustained responses to the same temporal sequence; scale bar is ΔF/F=2. **D.** Rank-ordered list of sustained response amplitude (average ΔF/F over the range t=10-17 sec) with threshold for defining a “strong” response (dashed line; ΔF/F ≥ 0.2). **E.** Activity versus time for 2 example neurons with transient plus periodic responses; scale bar is ΔF/F=1; dashed lines indicate baseline for each neuron. **F.** Activity versus time for 2 example neurons with maintained responses; scale bar is ΔF/F=0.5; dashed lines indicate baseline for each neuron.

Most sustained responses of individual neurons were periodic – namely, they exhibited one peak of calcium fluorescence during the entire temporal sequence. Sampling across different neurons, we found periodic responses with different phases (Fig. 6C). When we compiled results across multiple sequences and animals, we found a continuous distribution of amplitudes of the periodic response. This distribution was dominated by many small response amplitudes along with a few large amplitudes (Fig. 6D). Therefore, we set a criterion of an amplitude ΔF/F ≥ 0.2 to define a “strong” sustained response (corresponding to ∼6 spikes fired per sequence). By this definition, 200/5510 = 3.6% of all sustained responses were strong.

In addition, we observed three other kinds of less common neural responses. First, many neurons had both a strong periodic response and a transient elevation of firing when a new sequence was presented. We called these “transient plus periodic”; they comprised 77/5510 = 1.3% of all responses (Fig. 6E). Second, we found a second kind of sustained response that was roughly constant throughout the duration of the presentation of a given sequence, rather than periodically modulated (Fig. 6F). We called these “maintained” responses, and they could be readily distinguished from periodic sustained responses (Supplemental Fig. S8); they constituted 99/5510 = 1.8% of all responses. Third, some of the maintained responses increased in amplitude as a given temporal sequence was repeated; these “ramping maintained” responses were very rare, constituting 11/5510 = 0.2% of our measured responses (Fig. 6F *top*).

### The Two-Channel Population Code

Because each temporal sequence elicited sustained responses in different subsets of neurons, neural population activity has the capacity to encode the identity of the temporal sequence, rather than just the fact that a predictable sequence is occurring. Similarly, we wanted to know if the transient response was a generalized “alert” signal that would be invariant across different choices of the visual stimulus, or whether it depended on the stimulus in a fashion that allowed it to encode additional information about the identity of the temporal sequence. To follow up on these questions, we constructed a simple, cross-validated decoding algorithm that took single–trial neural responses and predicted which temporal sequence was ongoing (see Methods). This decoder had a linear form, with a set of weights for recognizing each of the 10 possible sequences. These weights were determined directly from a “training” fraction of the neural data (i.e. there was no learning or optimization over possible weights). Then, we applied the weights to a single trial of neural data to get a likelihood for each of the 10 sequences being present. Finally, the decoder's estimate was given by the sequence having the maximum likelihood. This decoder is a kind of multi-pattern linear classifier.

We wanted to separately study how transient versus sustained responses encoded sequence identity. To this end, we defined the transient response of each neuron, T, as its activity averaged over the first three cycles of a given temporal sequence, and the sustained response, S, as its activity averaged over the last three cycles (Fig. 7A). Then, we rank ordered all the sustained responses according to their amplitude, S. Direct inspection revealed that the most active sustained responses were large and highly stimulus selective (Fig. 7B). If we decoded using only the most active sustained cell for each temporal sequence, then we still got performance far above chance (Fig. 7D *open circles*). The second most active cell also gave substantial decoding performance, but much less than the most active cell. Performance then dropped off sharply as a function of rank order. Consequently, the cumulative decoding performance saturated quickly as a function of increasing population size (Fig. 7D *filled circles*). These results imply that the sustained responses employ a highly sparse code for sequence identity.

**Figure 7.**
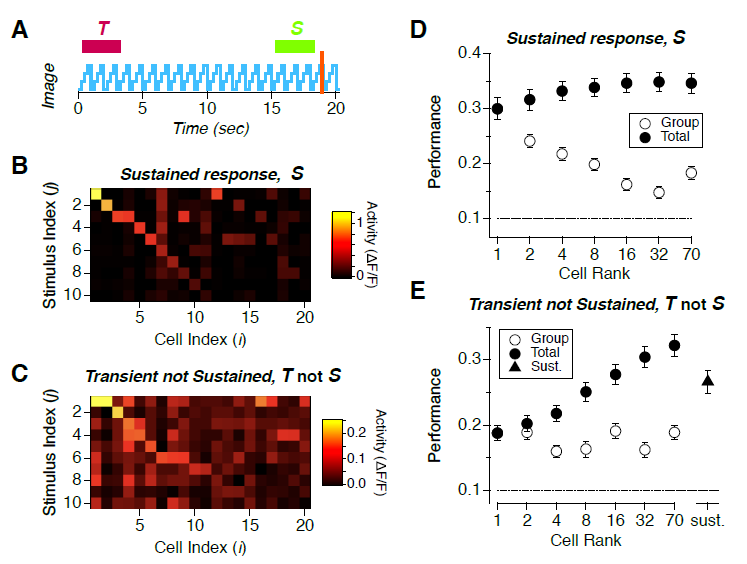
Decoding Population Activity. **A.** Image index versus time for a repeated temporal sequence with transient (T) and sustained (S) activity periods shown with colored bars (purple, green); novel image shown in orange. **B, C.** Matrix of activity (color scale) of cells, *i*, versus temporal sequences, *j*, for (sustained, transient not sustained activity). **D, E.** Decoding performance (fraction correct) for (sustained, transient not sustained) responses plotted versus cell rank for groups (open circles) and the cumulative population (filled circles); error bars are standard deviation across n=5 mice; chance performance shown with a dashed line; performance of the transient population in the sustained time window (triangle).

To study the transient code, we rank ordered responses according to the amplitude of the transient response, given the condition that the cell was not sustained, T not S. (Here, “not S” means that the sustained response amplitude was <0.2 ΔF/F.) Again, we could directly see that the most transient responses were stimulus selective (Fig. 7C). But in contrast to the sustained responses, transient responses were present for multiple temporal sequences and were weaker in absolute amplitude. When we decoded using subsets of cells with transient responses, we found that larger groups continued to allow significant decoding performance (Fig. 7E). At the same time, the cumulative decoding performance rose steadily as a function of rank order, reaching substantial performance with all of the purely transient responses. Furthermore, this performance was higher than for the the steady-state (sustained) responses of the same cells (Fig. 7E triangle; p < 0.001). This last observation demonstrates that the transient response encodes additional information about sequence identity not represented by the sustained responses. As a result, we can rule out the hypothesis that the transient response is merely a generic “alert” signal. Related to this, there was no elevated activity when the novel image was a blank, consistent with our finding that the transient response actively represents stimulus identity (Fig. S9). Together, these data imply that the transient response constitutes a dense code of relatively low amplitude and weakly tuned responses to represent sequence identity – very different from the sustained response.

We can visualize these results by constructing a schematic view of the transient and sustained population codes (Fig.8). The transient code has a majority of cells active but decays after several repetitions of the same sequence (Fig. 8A *i*). The sustained code has a sparse set of strong responses along with many weak responses; periodic responses have their own well-defined phase, and maintained responses are roughly constant in time (Fig. 8A *ii*). We can think of the transient and sustained responses as two channels of information within the neural population. However, these responses are not perfectly segregated in individual neurons – recall that some neurons are “transient plus periodic”. Thus, these two population codes are multiplexed to produce the activity of individual neurons (Fig. 8A iii). Then, when a different temporal sequence is ongoing, the population again contains these two channels of visual information with the same structure, but with different identities of which neurons exhibit transient and sustained responses (Fig. 8B).

**Figure 8.**
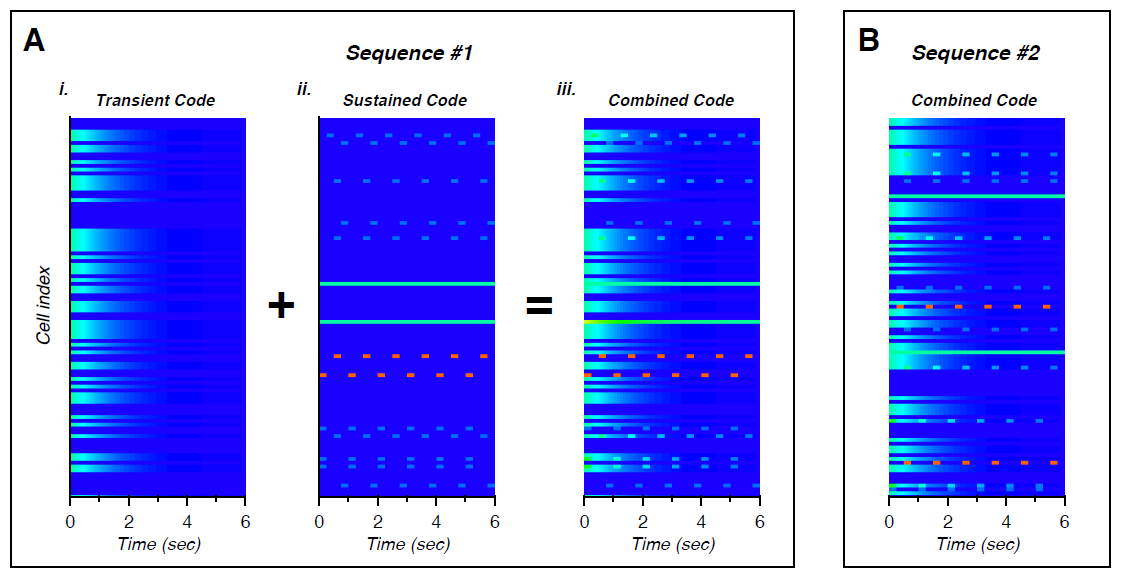
The Two-Channel Predictive Code. **A.** For a given sequence, the transient code (*i*) has many cells exhibiting moderate activity for the first few repetitions, the sustained code (*ii*) has a sparse set of cells with strong periodic modulation, another sparse set with maintained activity, and many cells with weak activity. These two codes are multiplexed together into the combination population activity (*iii*). **B.** For a different temporal sequence, the structure of the code remains the same, but the specific cells exhibiting each kind of activity are different.

## Discussion

Our results show that layer 2/3 of the primary visual cortex employs a population code with two channels of information: a dense transient population response that encodes novel stimuli and a sparse sustained population response that encodes repeated stimuli. This phenomenology seems quite general, in that we observed a transient response when we switched from one repeated stimulus to another, regardless of our choice of images or other stimulus parameters, such as the sequence length. Conceptually, a novelty response requires that the system embodies at least a statistical form of prediction about the regularity of the stimulus. In this sense, it is consistent with many theories of predictive coding that postulate that a neural system should only send surprising information up the sensory hierarchy (Barlow, 1961; Bastos et al., 2012; Rao and Ballard, 1999; Srinivasan et al., 1982). However, our results differ from these theories of predictive coding, because we also found a sustained population response that continues even for highly predictable stimuli.

Interestingly, a more recent theory of cortical computation has proposed that predictable stimuli need a positive representation and that this is the representation that should be fed forward up the cortical hierarchy (Heeger, 2017). The sustained response that we observed in layer 2/3 neurons has this basic property. Another property that has been proposed for sustained responses in a predictive code is that neural activity should encode a temporal sequence with a pointer-like representation (Hawkins and Blakeslee, 2004). This means that a neuron should fire once per temporal sequence rather than once per image of the sequence. This representation is advantageous for building a hierarchical predictive code, as the signals that are sent up the cortical hierarchy represent the entire temporal sequence as a single, composite sensory event. This then allows the next stage in the hierarchy to learn temporal sequences formed out of these composite events. Our data embody this property, in that strong periodic responses typically exhibited a single peak of activity during the temporal sequence. While this property could be interpreted as merely resulting from the sparseness of neural activity, the underlying mechanism does not change the significance of this representation for the neural code. Taken together, the two-channel predictive code combines properties posited by different theories of predictive coding, but which are not all present in any existing theory.

One notable property of the sustained population response was its sparseness – namely, that we observed only ∼2% of cells with a strong periodic response under our stimulus conditions. Sparseness is also a property that has long been as-sociated with efficient or predictive neural codes (Atick and Redlich, 1990; Attwell and Laughlin, 2001; Baddeley et al., 1997; Bastos et al., 2012; Bharioke and Chklovskii, 2015; Hawkins et al., 2009; Lennie, 2003; Olshausen and Field, 1996). Sparse responses have been reported in several studies (Froudarakis et al., 2014; Weliky et al., 2003; Willmore et al., 2011; Yen et al., 2007), and sparseness is enhanced by the extraclassical surround of V1 cells (Alink et al., 2010; Vinje and Gallant, 2000). For comparison, the lifetime sparseness of neurons during the *Repeated Sequence Experiment* was0.77 ± 0.10 (n = 551 neurons; see Methods). While a direct comparison of these numbers to those derived from extracellular spike trains would need to compensate for the temporal averaging of the calcium indicator, these numbers indicate a high but not unprecedented level of sparseness.

In the context of predictive coding, we should point out, however, that most of our observed sustained responses began during the first presentation of a new temporal sequence. Thus, they cannot result from the system learning the regularity of the new repeated stimulus, on the fly. Instead, their tuning properties may embody learning that occurred before we carried out our experiment. Thus, we prefer to say that these responses represent “familiar” rather than predicable stimuli. In this vein, neurons higher up in the visual pathway have also been observed to shift their tuning properties after extensive exposure to the same stimuli, such that most neurons decrease their firing to a familiar stimulus while a small fraction increase their firing (Li et al., 1993; Woloszyn and Sheinberg, 2012). Because sustained responses began during the first presentation of a new temporal sequence, it is natural to think of sustained responses as being driven significantly from the feedforward pathway, including connections from layer 4. Thus, the sustained response can embody features found in feedforward models of cortical function (Heeger et al., 1996; Hubel and Wiesel, 1962; Rust et al., 2005).

The transient response is closely related to the phenomenon of stimulus-specific adaptation (SSA), which proposes that the response of a cortical neuron to an infrequent stimulus is larger than the response to the same stimulus when it is more common (Hamm and Yuste, 2016; Natan et al., 2015; Ulanovsky et al., 2003; Yaron et al., 2012). For instance, the novel image that we showed in the *Novelty Experiment* was always a unique image, which is maximally infrequent, and hence should generate a very large response according to the logic of SSA. Similarly, the presentation of new stimulus sequences in the *Variable Repetition* or *Sequence Length Experiments* should generate a large initial response due to SSA. The comparison also applies to *Repeated Sequence Experiment*, where a handful of temporal sequences were repeated several times: If the time scale over which stimulus frequency is effectively measured is short compared to the duration of a stimulus block (20 sec), then the stimulus sequence that begins in a new block should also be an infrequent stimulus.

However, the overall picture encompassed by the idea of a two-channel predictive code differs from that entailed by SSA. In SSA, every neuron has an adaptation index that characterizes its degree of adaptation, and it is not clear if a neuron will respond to a rare stimulus that is outside of its classical tuning curve. In our experiments, we found that a given neuron could have a transient response or not depending on the identity of the stimulus. In the language of SSA, this would correspond to a different adaptation index for all pairs of stimuli tested. Furthermore, we observed many neurons to have a purely transient response, meaning that they initially responded to a stimulus that was outside of their classical tuning curve. These differences could very well follow from different approaches to experimental design. In particular, experiments probing SSA having largely taken a single cell point-of-view, while the experiments here have taken a population coding point-of-view. SSA exhibits a variety of timescales (Ulanovsky et al., 2004), perhaps matching the flexible dynamics of the transient response (Fig. 5).

One possibility is that the transient response results from a form of contrast adaptation (Dhruv and Carandini, 2014; Kohn, 2007; Muller et al., 1999). This is because the effective local contrast on a given neuron's receptive field can vary between images, even though the global contrast is held fixed. However, the phenomenology of contrast adaptation is somewhat different. When local contrast increases, neurons will desensitize and reduce their activity, but when contrast decreases, neurons will correspondingly resensitize and increase their activity during adaption. The latter process is almost entirely absent in our data. Furthermore, contrast adaptation typically leads to a substantial steady-state level of activity, while we observe the transient response to decay to essentially zero steady-state activity for most cells. Therefore, we do not believe that the transient response should be interpreted as a simple form of contrast adaptation.

Perhaps the transient response can be instead viewed as a form of adaptation to a very general class of contingencies? This idea has been proposed for the visual cortex (Carandini et al., 1997; Movshon and Lennie, 1979), and many weaker forms of such adaptation have been observed in the retina (Hosoya et al., 2005; Smirnakis et al., 1997). But again, these more general forms of adaptation result in substantial steady-state activity. Furthermore, the transient response exhibits dynamics that are both faster than these higher order forms of adaptation and more flexible, in that the dynamics scale with sequence length. Thus, while the transient response can certainly be interpreted as a form of adaptation, it is at the very least a “stronger” form than has been previously reported.

The transient response also resembles the mismatch negativity (MMN). Originally described as a negative deflection in EEG voltage in response to an unexpected or oddball auditory stimulus (Naatanen et al., 1982), the term has come to encompass analogous phenomena in other sensory pathways and measured with other brain imaging techniques, like MEG or field potentials (Naatanen et al., 2007). Common to all experimental paradigms is that the MMN is manifested in the average activity of populations of neurons. When we construct the population average of neural activity, we see elevated activity after a novel visual stimulus. We believe that this elevated population activity is analogous to the MMN. But our cellular resolution imaging reveals that this is not the entire story: namely, that there is also a sparse subset of sustained responses to repeated stimuli.

How is the transient response generated? Because of its similarity with stimulus-specific adaptation, we should compare it with the mechanisms underlying SSA. Geffen et al. showed that two classes of inhibitory interneurons in L2/3 shape and enhance SSA (Natan et al., 2015). But at the same time, the phenomenon was already present in the current source density in L4, suggesting that synaptic depression in the MGN → L4 synapse may play an important role. While this mechanism must make some contribution, it is unlikely to be sufficient to explain the transient response for two reasons. First, the time scale is too short. Varela et al. showed that the dominant form of synaptic depression in L4 → L2/3 synapses in V1 recovered with a time constant of ∼400 ms (Varela et al., 1997). Similar depression has been observed in LGN → L4 synapses (Boudreau and Ferster, 2005; Chance et al., 1998; Gil et al., 1997). This form of synaptic depression would largely recover after each image presented in our experiments, preventing a build-up of depression that could account for the transient response. Second, the flexibility of dynamics as a function of sequence length (Fig. 5) cannot be described by a biophysical process with a fixed time constant, like synaptic depression.

Flexible dynamics and long timescales might both emerge, in part, from cell intrinsic mechanisms. Lundstrom et al. injected white noise currents into cortical pyramidal cells and showed that the spiking response exhibited a broad range of timescales described as a process of fractional differentiation (Lundstrom et al., 2008). Mechanistically, this process can emerge from the action of slow conductances having a range of time constants. Flexible dynamics has also been observed in retinal adaptation, where the time constant of adaptation scaled with the time over which stimulus statistics were changed (Wark et al., 2009). This phenomenon closely resembles how the dynamics of the transient response scale proportionally to the sequence length, suggesting that retinal adaptation might be making a contribution. Yet neither of these mechanisms is likely to provide a complete description of the transient response, as they both cause adaptation to a lower steady-state response, while the transient response decays to zero baseline. One final possibility is that the retina recognizes periodic sequence violations via the omitted stimulus response (Schwartz et al., 2007a). However, we specifically designed each image to last for 250 ms or more, which is beyond the memory of retinal periodic pattern detection.

Central to many theories of predictive coding is the idea that inhibitory neurons encode predictable stimuli and feed that inhibition onto excitatory principle neurons to cancel their response (Bastos et al., 2012; Bharioke and Chklovskii, 2015; Srinivasan et al., 1982). One salient example of this mechanism is the cancellation of predictable stimuli in cerebellum-like structure of weakly electric fish (Bell et al., 2008). One requirement for this mechanism of active cancellation is that some neurons would need to have their activity ramp up in a fashion complementary to the decay of the transient response. In this vein, we did see some neurons with ramping maintained responses, but these cells were very rare. However, it is important to point out that our transgenic mouse line only expressed calcium indicator in excitatory cells (Dana et al., 2014), so it is still possible that one or more classes of inhibitory interneuron have ramping responses. Thus, it will be interesting to record specifically from inhibitory cells in subsequent studies. Of course, any inhibitory cancellation mechanism must simultaneously leave the sustained responses unaffected – a property that serves as an important constraint on the underlying circuit mechanisms.

A recently proposed model of temporal sequence learning in layer 2/3 agrees with many qualitative features of our data (Hawkins and Ahmad, 2016). Once a temporal sequence has been learned, the occurrence of the initial elements of the sequence put pyramidal cells into a “predictive” state. When the prediction comes true, such a neuron fires early and helps to generate inhibition that silences neighboring neurons. This firing continues for repeated stimuli, as we observe for the sustained response. If instead the prediction is violated, then a larger group of neurons fire together before being silenced by inhibition. Thus, there is a larger network response to a stimulus event that violates an ongoing prediction, as we observe for the transient response. However, a detailed comparison of the properties of this model and our data has not been carried out.

In its broad form, the two-channel predictive code that we have observed in the primary visual cortex shares many similarities with the code found in the retina (Berry and Schwartz, 2011). The population code of retinal ganglion cells separately encodes information about temporal sequences of sensory events that are repeated versus novel into two different information channels (Schwartz et al., 2007a; Schwartz et al., 2007b). As in V1, this separation is not observed at the single cell level – namely, there are no “cell types” specialized for repeated versus novel stimuli. Instead, individual neurons participate in both information channels, and the two channels themselves are only defined at the population level. This broad similarity suggests that this might be a coding strategy employed by neural populations in many brain areas.

However, there are also many detailed differences between these predictive codes in the retina versus V1. In particular, the retina has only a limited capacity to recognize repeated temporal sequences. For the case of periodic sequences of flashes, the retina only recognizes violations for periods of up to ∼200 ms; for flashes separated by longer time intervals, the retina effectively treats the next flash as novel (Schwartz and Berry, 2008; Schwartz et al., 2007a). In contrast, V1 exhibits predictive coding for temporal sequences of up to 3.6 sec in duration (Fig. 5), as one might expect due to the extensive recurrent excitatory circuitry in the neocortex. Thus, the neocortex may possess the ability to recognize a far more general class of longer and more complex temporal sequences.

## Methods

### Animal Surgery and Husbandry

All experiments were performed according to the Guide for the Care and Use of Laboratory, and procedures were approved by Princeton University's Animal Care and Use Committee. We used animals from transgenic mouse line GP5.3 supplied by Janelia Research Institute, which expresses GCaMP6f under the Thy1 promoter (Chen et al., 2013; Dana et al., 2014). Calcium activity was imaged through a chronic cranial window. The implantation of the cranial window followed steps similar to (Dombeck et al., 2010). In short, anesthesia was induced in mice with 2.5% isoflurane, and then maintained at 1.5% during the surgery. A 5 mm round craniotomy was drilled with a dental drill centered around 2 mm posterior and 1.75 mm lateral from bregma. The dura was left intact. The hole was then covered with a canula consisting of a 5 mm round microscopy cover glass glued to a metal ring. The metal ring was glued to the bone around the craniotomy with vetbond. Then the skull around the craniotomy was covered with metabond. A titanium head plate was then attached to the skull and secured with metabond. After the surgery, mice were provided with pain management and allowed to recover for several days before any recordings. Animals were placed on a reversed night-day cycle.

### Quantifying Animal Behavior

Mice were placed on a water restriction schedule with 1.5 ml per day. Mice were then head fixed and allowed to run on a Styrofoam wheel. A lick port was positioned in front of the animal's mouth, which could make water available via a solenoid valve. Temporal sequences were presented on a computer monitor with the design of the *Novelty Experiment*. Each image (Gabor or grating) was presented for 200 ms. Initially sequences were formed from S = 2 or 3 images, and the final image of a sequence was stochastically replaced with a novel image at a rate of once per 6 sequences. During the initial, 3-day shaping period, water was made available at the lick port after 50% of all novel images. In doing so, a solenoid valve made a clicking noise that the animal could hear. Animals readily learned to obtain water in these “automatic” trials. On the other 50% of violation trials, which we denote as “test” trials, water was only available after the animal started licking (and within a time window 0.5 – 1.5 s after the violation).

After the shaping period, water was made automatically available after only 20% of all novel images. We quantified performance on the other 80% of novel images by measuring the rate of lick onsets relative to the time at which the novel image was presented. A lick onset was defined as a lick that occurred more than 800 ms after the previous lick. Once animals reached a performance criterion (significance of peak lick onset rate p < 1e-10), the sequence length was increased by adding 1 new image to the old sequence and training continued.

### One-Photon Ca++ Fluorescence Imaging and Mapping of Visual Areas

Mouse visual cortex consists of several discrete areas. In order to ensure that all our recordings were performed in the primary visual cortex (V1), we generated a map of the visual cortex that showed the boundaries of the visual areas with respect to the vasculature. We then used the vasculature as landmarks in order to select a recording area for our 2-photon imaging that lay safely within the boundaries of V1. Visual area boundaries were mapped with drifting bars presented on an LCD screen as described in (Marshel et al., 2011). In short, the mouse was placed on an air suspended styrofoam ball and fixed to a post using its head plate. An LCD screen was placed to the right of the awake mouse at a distance of 15 cm and an angle of 10 degrees in order to be parallel with right eye, covering a visual field reaching form 150 degrees vertical to 145 degrees horizontal. A bar, consisting of blinking checkers was then slowly moved many times across the field of view in all four cardinal directions. During the presentation of this stimulus, the cranial window was imaged at 30 Hz with a fluorescent microscope with a field of view covering the whole 5 mm diameter round cover glass. The resulting video of bulk fluorescent activity was then analyzed to find boundaries at which the spatial map reversed. These boundaries were used to segment visual areas. Finally, anatomical landmarks were used to identify the primary visual cortex (V1) using an algorithm described in (Garrett et al., 2014).

### Two-Photon Ca++ Fluorescence Imaging

Mice were fixed to a post using their head plates and were free to run on a large, air-suspended styrofoam ball. The rotational speed of the ball in both axes was acquired with an optical mouse and synced with the neural data. A custom made 2-photon microscope (NA 0.8, 40x water immersion objective) was used to image a 400 x 400 micrometer area within the cranial window. The setup is described in detail in (Dombeck et al., 2010). GCaMP6f was excited at 920 nm with a Titanium sapphire laser (140 fs pulses at 80 MHz). Fluorescence was detected with a photomultiplier tube. Laser path and image acquisition was performed with the help of ScanImage 5.0. We used the areal map established with the one-photon fluorescence rig and used the vasculature as landmarks for finding V1. Two-photon laser light (60 mW) was then focused at the highest cell density at around 200 – 300 µm cortical depth, corresponding to layer 2/3. Images were acquired with 512 scan lines at 30 Hz, resulting in a movie of calcium fluorescence with 512 x 512 pixel frames (covering a 410 x 410 µm field of view).

Visual stimuli were presented to the mouse via a projection system. A digital projector image was spread via an angular amplification mirror to fill a toroidal projection screen. The field of view from the point of the mouse reached from roughly -20 to +70 degrees vertically and -130 to +130 degrees horizontally. Neural data and stimuli were synchronized through analog signals recorded in Clampex. In a subset of experiments, the contralateral eye was filmed during the stimulus presentations at 60 Hz with an infrared video camera. From this this data, pupil diameter, pupil position, and eye movements and blinks were extracted with a custom code written in Matlab (see Supplementary Figures S5, S6).

### Design of Visual Stimuli

We formed spatial images with either a random mixture of Gabor functions (Gabor) or a static, square-wave grating (grating). To form Gabor images, 100 Gabor functions were randomly chosen with: 100% contrast, either ON- or OFF-polarity, random location, random orientation, and random phase. We picked a spatial extend for each Gabor randomly from 10°-20° matching receptive field sizes in V1 neurons in mice. Gabor functions were linearly superimposed with saturation at 100% contrast. The overall light level of images therefore varied by standard deviation of 1.2% of the mean. Grating images had a spatial period of 20°, random phase, and random orientation.

Temporal sequences were composed of multiple, randomly selected spatial images. Images and sequences were presented without blank frames in between. The specific composition of temporal sequences varied in each of 4 experiments:

#### Novelty Experiment

200 ms per image, 5 images per sequence, repeated for ∼10 min. The final image of the sequence was replaced with a novel image at a rate of once per 6 sec. Two 10 min blocks with different temporal sequences were used.

#### Variable Repetition Experiment

300 ms per frame, 3 images per sequence. Sequences were formed into adaptation blocks with L = [3, 4, 6, 11, 21, 41] total cycles. The final image of the second-to-last sequence was replaced with a novel image. Each adaptation block had a new choice of temporal sequence and a new choice of L. Adaptation blocks were presented without blank frames in between for ∼60 min.

#### Variable Sequence Length Experiment

300 ms per frame, S = [3, 6, 9, 12] images per sequence, and 21 cycles per block. The final image of the second-to-last sequence was replaced with a novel image. Each adaptation block had a new choice of temporal sequence and a new choice of S. Adaptation blocks were presented without blank frames in between for ∼60 min.

#### Repeated Sequence Experiment

300 ms per image, 3 images per sequence, 21 cycles per adaptation block. The final image of the second-to-last sequence was replaced with a novel image. Each adaptation block had a random choice of one out 10 possible temporal sequences. Adaptation blocks were presented without blank frames in between for ∼57 min, such that each temporal sequence was presented for a total of 18 times.

### Data Preprocessing

Mouse movements could cause a slight frame-to-frame shifts in the recorded field of view. We therefore aligned all frames to a common template, constructed. by averaging 1000 individual frames. Then, individual frames were shifted to maximize cross correlation with regards to the template (Dombeck et al., 2010).

Regions of interest (ROIs) corresponding to neurons were identified based on time-averaged images (Apthorpe et al., 2016). Both obviously active and seemingly inactive ROIs were selected for processing. A typical field of view contained about 100 ROIs. Pixel values belonging to a given ROI were averaged on a frame-by-frame basis and converted to relative fluorescence, ΔF/F. The baseline, F, of the fluorescence signal was estimated by binning the data within 5 minute windows and taking the mode of the corresponding distributions.

### Quantifying the Novelty Response

For each ROI, we subtracted the peri-stimulus time histogram (PSTH) triggered on the start of each repeated temporal sequence from the PSTH triggered on the time of a novel image. This gave us the excess response to a novel image, *ΔR*. To estimate statistical significance, we calculated the average neural activity (ΔF/F) in a time window 300–500 ms following each novel image and subtracted the baseline neural activity in a time window 1000 ms before the novel image, giving *ΔR*_*i*_ for each instance of a novel image *i*. We calculated the standard error of the mean over all instances of a novel image, *δR*, and used this to calculate the z-score, *z* = Δ*R*/ δ*R*. From the z-score, we calculated the probability that a positive excess response could occur by change (p-value).

### Excess Spikes Evoked by a Novel Image

The density of neurons in layers II and III of the mouse primary visual cortex is roughly 150,000 neurons/mm3 (Schuz and Palm, 1989). This, together with a rough estimate of the thickness of layer 2/3 V1 in mouse of 250 µm taken from (Smith et al., 2009) and an estimated area of roughly 4 mm2 (Marshel et al., 2011), gives 150,000 neurons in V1 layer 2/3 per hemisphere. Given our estimate of 0.5 extra spikes per neuron (Supplemental Fig. S1), this amounts to roughly 150,000 extra spikes within a short time window across the two hemispheres of V1.

### Decoding Analysis

In order to use population neural activity to estimate which temporal sequence was ongoing, we constructed linear, multi-pattern decoders. We denote the neural activity of cell *i* at time *t* during trial *k* of sequence *j* as: *r*_*ijk*_(*t*). This neural activity was evaluated over different temporal windows for the transient versus sustained responses (see main text). The linear decoder convolves the neural activity measured on a single trial *k* with a weight vector, *w*_*ij*_, to get a likelihood for sequence:

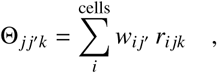

where *j* denotes the real, ongoing sequence and *j'* denotes all possible sequences. The estimated sequence is given by the maximum likelihood:

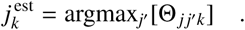

Then, the decoding performance is given by the fraction of trials *k* that get the right answer,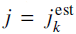. Given a test trial *k*, the weights were determined in a cross-validated fashion from the training data (all *k'* ≠ *k*) with no optimization:

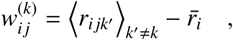

*where 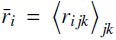* is the average activity of neuron *i* over all trials and sequences. Notice that the weights differ for each test trial, because of the cross-validation procedure. In all cases, when we rank ordered the activity of neurons, this ranking was carried out separately for each temporal sequence. In implementing our decoder, we pooled cells from across different mice and ignored noise correlation.

### Lifetime Sparseness

Following previous studies, we defined the lifetime sparseness of a neuron *i* as:

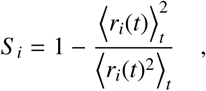

where the average is over all time bins. For this calculation, we averaged over all trials of the same stimulus to get a PSTH for each cell, *r*_*i*_(*t*). We performed this calculation for the *Repeated Sequence Experiment*, in which each neuron was presented with 10 different temporal sequences. We concatenated the PSTHs across all 10 temporal sequences and then computed the lifetime sparseness of each neuron.

## Acknowledgements

The authors thank Winfred Denk, Jeff Gauthier, Rava Azeredo da Silveira, Adrienne Fairhall, and Carlos Brody for useful discussions. The authors also thank Ilana Witten, Chris Honey, and Subutai Ahmad for helpful comments on the manuscript. This project was funded by grant R21EY026754 from the National Eye Institute as well as support from the Princeton Neuroscience Institute.

